# DASH-CAM: Dynamic Approximate SearcH Content Addressable Memory for genome classification

**DOI:** 10.1101/2023.09.29.560142

**Authors:** Zuher Jahshan, Itay Merlin, Esteban Garzón, Leonid Yavits

## Abstract

We propose a novel dynamic storage-based approximate search content addressable memory (DASH-CAM) for computational genomics applications, particularly for identification and classification of viral pathogens of epidemic significance. DASH-CAM provides 5.5× better density compared to state-of-the-art SRAM-based approximate search CAM. This allows using DASH-CAM as a portable classifier that can be applied to pathogen surveillance in low-quality field settings during pandemics, as well as to pathogen diagnostics at points of care. DASH-CAM approximate search capabilities allow a high level of flexibility when dealing with a variety of industrial sequencers with different error profiles. DASH-CAM achieves up to 30% and 20% higher *F*_1_ score when classifying DNA reads with 10% error rate, compared to state-of-the-art DNA classification tools MetaCache-GPU and Kraken2 respectively. Simulated at 1GHz, DASH-CAM provides 1, 178× and 1, 040× average speedup over MetaCache-GPU and Kraken2 respectively.

**CCS CONCEPTS:** •**Hardware** → **Bio-embedded electronics**.

## 1 INTRODUCTION

Content-addressable memories (CAMs) offer outstanding performance in applications where high-speed searching is critical [46, 55]. In addition to well-studied applications, such as network routers, digital signal processing, analytics, and reconfigurable computing [30, 46], CAMs can be used in a variety of emerging compare-intensive big data workloads [34], machine learning applications [2, 23], and bioinformatics [27, 29, 53].

While CAMs are typically designed to find exact matches, approximate or similarity search is increasingly required by many popular contemporary data-intensive applications, such as data analytics, machine learning, deep learning, and computational genomics [26, 28, 29]. The latter gained keen interest of the research community due to the exponential growth of the sequenced DNA data volumes in recent years [52]. It is an active research field and a basis for a variety of applications, such as genomic surveillance, antimicrobial resistance diagnostics, environmental ecosystems monitoring, sustainable agriculture, and personalized healthcare [3, 18, 51, 59].

In approximate search, if the difference between a stored pattern and the query pattern is below a certain predefined threshold, such stored pattern is considered a “match”. While in some cases, the difference is limited to replacements (i.e., where a data element is replaced by another), in applications involving text and sequenced DNA data, there are two additional types of difference: insertions (where a data element is inserted into an otherwise identical sequence of data elements), and deletions (where a data element is deleted from a sequence), collectively called indels. Replacements, insertions and deletions are typically referred to as edits.

A variety of CAM designs support approximate search. These solutions target Hamming distance tolerance, typically limited to only a few bits [47]. HD-CAM [15, 16] proposed an approximate search CAM design that can tolerate very large Hamming distance (50% of the data wordlength and above). However, HD-CAM is SRAM based and hence its scalability is limited. In consequence, in applications like genome classification, both the number of genome classes and genome sizes are limited due to said lack of scalability.

To alleviate this fundamental limitation, we propose a dynamic approximate search content addressable memory (DASH-CAM), which is capable of tolerating a user - configurable Hamming distance while providing 5.5× density compared to HD-CAM. DASH- CAM targets the high speed approximate search in applications such as computational genomics, particularly in detection and identification of pathogens.

Traditionally, pathogen detection relies upon the identification of pre-established markers of a particular disease [50]. However, pathogen detection increasingly deal with metagenomic samples, for example sourced from wastewater [5]. In particular, genome identification is performed as follows (Fig. 1): (1) a sample potentially containing DNA of multiple organisms is obtained and prepared; (2) the sample is sequenced; the sequencer outputs DNA reads (potentially sourced from different organisms); DNA sequencers are prone to sequencing errors, such as indels and replacements; (3) DNA reads (soiled with sequencing errors) are processed by a metagenomic classification application that potentially associates each DNA read with a certain species in its database. A variety of DNA classifiers have been proposed, including Kraken2 [54], which classifies DNA by exact matching of DNA read fragments (called *k*-mers) against an existing DNA database, e.g., a collection of pathogen DNA. Since DNA reads typically contain sequencing errors, a certain fraction of query *k*-mers would not hit in the database, thus limiting the sensitivity of conventional DNA classifiers. The operating principle of DASH-CAM is based on the observation that the speed of matchline (a signal by which a match or a mismatch is signalled in a CAM row) discharge is proportional to Hamming distance between a query pattern and the data pattern stored in a CAM row. If the matchline voltage is above a certain voltage threshold at the sampling time, we consider it a match. Conversely, if the matchline voltage level is below said threshold at the sampling moment, it means that the query pattern misses in DASH-CAM.

**Figure 1:**
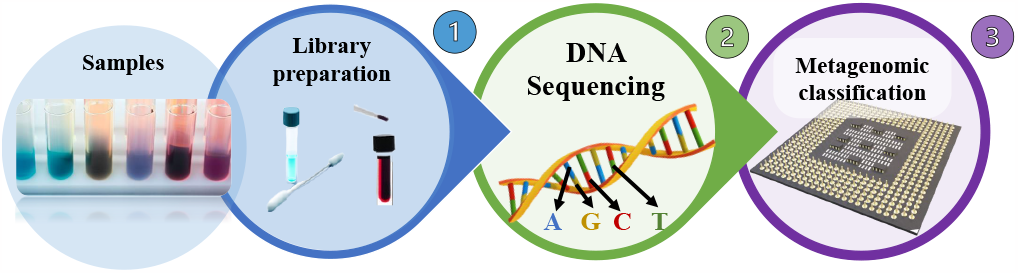
DNA classification pipeline comprising library preparation, DNA sequencing and computational genomics based detection and classification.

Our paper makes the following main contributions:

- To our knowledge, DASH-CAM is the first dynamic approximate search CAM that can tolerate user - programmable Hamming distance, designed for genome classification application.
- DASH-CAM based genome classifier design uses one-hot encoding of DNA bases to mitigate the retention time variation and potential data loss (which in DASH-CAM does not adversely affect the classification sensitivity).
- Overhead-free refresh: DASH-CAM refresh is performed in parallel with the main DASH-CAM operation, thus causing no performance degradation.
- DASH-CAM is evaluated as a part of pathogen detection and identification platform, to be used for example for pathogen transmission and mutation tracking during viral pandemics.

## 2. BACKGROUND AND PRIOR ART

### 2.1 Content-Addressable Memory

Fig. 2 shows the architecture of a conventional CMOS content-addressable memory (CAM) array comprised of *n* columns and *m* rows [46]. The CAM performs a comparison between the query data pattern stored in the search data register, and the information contained within the six-transistor static random access memory (6T-SRAM) bitcells. A matchline (ML) is shared between bitcells of an *n*-bit word and also fed into a sense amplifier (SA). The searchline (SL), denoted SL and 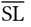, are shared across all rows of the CAM array. Read and write operations within the CAM array are executed in a similar manner as in conventional 6T-SRAM. Access to a bitcell is facilitated by enabling the word line (WL) for the corresponding row and precharging or asserting (complementary values) the SLs for read or write operations, respectively. A search (compare) operation is performed simultaneously across the entire array during a single clock cycle by asserting the query data on the SLs. The matchline sense amplifiers (MLSAs) evaluate the state of the matchlines at the end of the comparison cycle and signal a match or mismatch.

**Figure 2:**
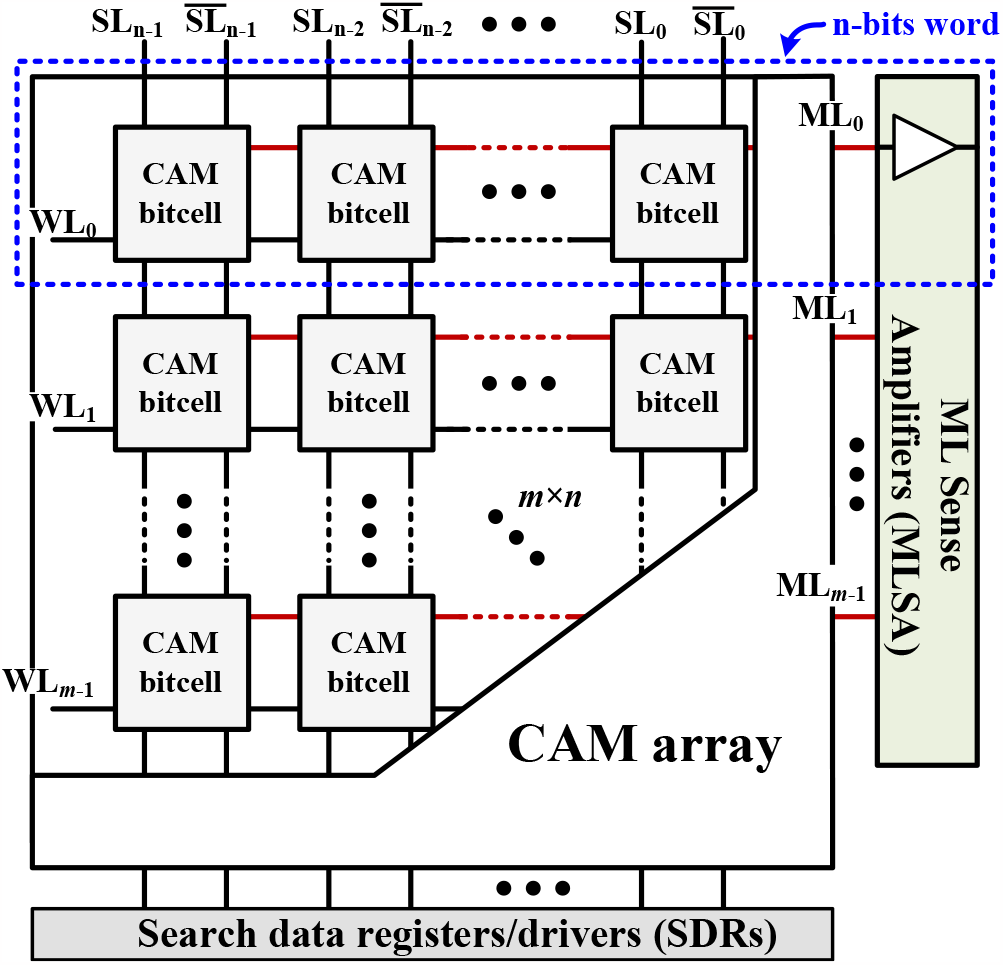
Conventional content-addressable memory (CAM) array.

Several CAM designs utilize emerging nonvolatile memories, such as STT-MRAM [17, 19], FeFET [58] and ReRAM [56, 57]. While such designs provide considerably higher density compared to SRAM, they suffer from high power consumption and limited endurance during write operations. They are also slower and more expensive to manufacture than CMOS solutions. Additionally, resistive memories are highly susceptible to process variation. DASH-CAM combines high-speed, practically unlimited write endurance, low-power consumption, reliability and low cost of conventional CMOS with the density of resistive memories.

### 2.2 Approximate Search Content-Addressable Memory

In recent years, many ternary and binary NOR- and NAND-based CAM bitcell designs have been proposed, including both CMOS-based [11] and emerging-memory-based [25] solutions. Several CAM designs offer soft-error tolerance using error correction coding (which requires memory redundancy) and replacing the matchline (ML) sense amplifier with an analog comparator [33, 45]. Such designs typically only tolerate a limited Hamming distance (1–4 bits). Another class of approximate search CAMs uses locality sensitive hashing of stored data and query patterns [39, 48]. While such schemes potentially tolerate large Hamming distances, they require hashing of data prior to storage and search. Additionally, large Hamming distance does not always result in low similarity of hashed data sketches [36], which leads to false positive results and hence limited precision.

Several emerging memory (memristor crossbar) based designs for Hamming distance approximation have also been proposed [53, 60].

Some of the approximate search CAM designs use timing (i.e., the matching score signal delay, or the speed of the ML discharge) as a measure of Hamming distance. HD-CAM [15], a Hamming distance-tolerant CAM, uses the combination of the voltage, controlling the speed of the ML discharge, and the sense amplifier reference voltage to define the Hamming distance threshold. HD-CAM is capable of tolerating very large Hamming distances. However, HD-CAM requires 3 bitcells to code a single DNA base, therefore the cost of storing one DNA base is 30 transistors (of both PMOS and NMOS types). A Hamming distance search CAM, where the matching score signal is delayed every time a bit mismatch occurs, is proposed in [8]. In the approximate search enabled CAM for energy efficient GPUs, proposed in [47], a small Hamming distance (≤ 2 bits) is tolerated through meticulous timing of the ML discharge. In [24], Hamming distance of up to 4 bits is tolerated by using delay lines at the clock inputs of four separate sense amplifiers on each ML. Tunable sampling time techniques require very precise device and circuit sizing, while achieving limited sensitivity and precision (due to false mismatches and multiple false matches [47]).

Recently proposed EDAM [20] is a CMOS edit distance-tolerant content addressable memory for approximate search. However, EDAM cell is very large (42 transistors) and hence scaling EDAM to support a large genome database is challenging. Additionally, EDAM design requires cross-connectivity among neighboring memory columns (to enable comparison with the left and right neighbors) which may render it wire-bound, adversely affecting density and timing.

### 2.3 Gain Cell eDRAM

The Gain Cell (GC) eDRAM is a type of embedded memory that uses two to four transistors, compared to the six transistors required for SRAM. GC-eDRAM designs use parasitic capacitance to store their memory state as charge and therefore require a periodic refresh. GC-eDRAM typically provides two-ported functionality (i.e., features separate read and write access ports).

Fig. 3 shows the 2T GC-eDRAM bitcell, comprising only nMOS transistors (NW and NR) that serve as write and read access ports. To perform a write operation, the write wordline (WWL) is asserted with a boosted positive voltage (i.e., WWL = V_BOOST_), to compensate for the threshold voltage drop across NW. This transfers the data (‘1’ or ‘0’) asserted on the write bitline (WBL) to the storage node capacitance (C_Q_), mainly consisting of the gate capacitance of NR and the junction capacitance of NW. For the read operation, the read bitline (RBL) is first precharged to *V*_DD_. When the read wordline (RWL) is asserted (RWL = 0 V), the RBL remains at *V*_DD_ if ‘0’ is stored in the node Q, or is discharged to ground, if ‘1’ is stored (Q = *V*_DD_). Finally, the output data is obtained by comparing the RBL to a reference voltage using a sense amplifier.

**Figure 3:**
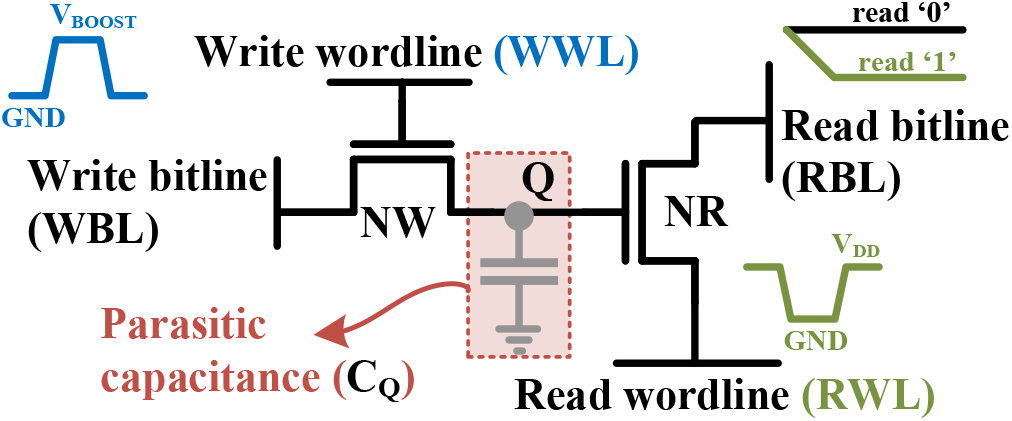
2T all-nMOS gain-cell implementation including the schematic waveforms.

### 2.4 Genome classification and profiling

DNA is composed of four nucleotides: Adenine (A), Guanine (G), Cytosine (C), and Thymine (T), which are frequently referred to as DNA basepairs or bases. Accordingly, a DNA data element is a DNA base that can have one of four values (A, G, C and T). DNA sequencing is the process of determining the bases of a DNA chain. Contemporary high-throughput DNA sequencers can sequence multiple DNA samples in parallel [22].

The goal of DNA classifier is to find what organism a DNA sequence (or a single DNA read) belongs to, taking into account that it possibly originates from a metagenomic sample (i.e., a sample that contains DNA of many different species). Several probabilistic classifiers have been proposed, such as interpolated Markov model based Phymm and PhymmBL [7], BLAST-based models [4], MEGAN [12] and METAPHYLER [35], naive Bayesian classifier NBC [49], and others. These classification tools are sensitive but relatively slow. For example, DNA classification using Smith-Waterman like dynamic programming would have the complexity ranging from *O* (*m* · *n*^2)^ (the best case) to *O* (*m*^2^ ·*n*^2)^ (the worst case), where *m* is the number of DNA reads and *n* is the read length.

Recently, convolutional neural network based classification solutions that differentiate between coronavirus (SARS-CoV-2) and other organisms have been proposed [1, 9].

A class of DNA classifiers uses exact pattern matching. Such classifiers include CLARK [44], CLARK-S [43], and Kraken2 [54]. MetaCache-GPU is a locality sensitive hashing based metagenomic classifier optimized for GPU [32]. These classifiers exhibit significantly higher speed at the cost of limited sensitivity. One reason for the reduction in sensitivity is sequencing errors that are inherently present in DNA reads. These sequencing errors manifest in replacing bases in DNA reads with incorrect ones, deleting bases, or inserting redundant bases. As a result, DNA read fragments (*k*-mers) that otherwise should have matched in the classification database, end up being unclassified and discarded.

Approximate search CAMs alleviate this fundamental flaw by tolerating to some extent the sequencing errors. When comparing a sequence *s*_1_ against a set of sequences *S* contained in an approximate search CAM, every *s*_2_ ∈ *S* such that *Hamming Distance (s*_1_, *s*_2_) ≤ *t* for some threshold *t*, will match. To endure such sequencing errors, the tolerance threshold *t* needs to be relatively high. This increases the sensitivity of classification (by lowering the false negative rate), but at the same time, reduces its precision (by elevating the false positive rate).

State-of-the-art classifiers include hardware-accelerated solutions as well. SquiggleFilter [13] and RawHash [14] are virus detection and classification solutions that analyze the raw output (raw squiggles) of the ONT MinION sequencer [41]. On the opposite side of complexity spectrum, GenSLMs[61] applies large language models to identification and classification of viral variants using supercomputers such as Polaris at the Argonne Leadership Computing Facility and Selene at NVIDIA, as well as Cerebras CS-2 wafer-scale cluster.

In this work, we present a fast, highly sensitive and precise approximate matching-based DNA detection and identification solution, implemented by DASH-CAM.

## 3 DYNAMIC APPROXIMATE SEARCH CAM DESIGN

We solve the lack of scaling of state-of-the-art solutions such as EDAM [20] by introducing DASH-CAM, whose design is presented in Fig. 4. To significantly increase the memory density, we replace a CMOS SRAM bitcell by a two-transistor dynamic memory cell. DASH-CAM stores the DNA bases in one-hot encoding format, such that three SRAM bitcells of HD-CAM [15] are replaced by four dynamic memory cells. Furthermore, by adding just four NMOS transistors, we implement the comparison logic that requires several logic gates in state-of-the-art designs.

**Figure 4:**
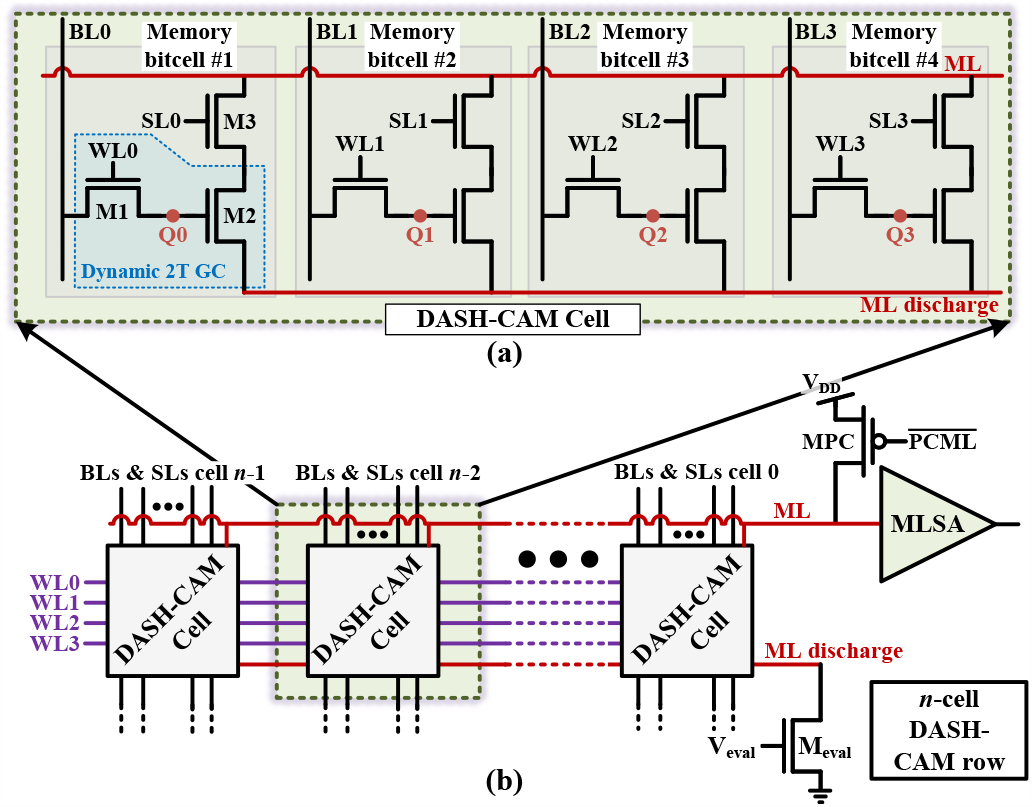
Dynamic Approximate SearcH CAM (DASH-CAM): (a) cell, (b) row.

### 3.1 DASH-CAM array design and operation

Fig. 4(a) presents the 12T DASH-CAM cell, comprising four 2T (M1-M2) gain cells and additional four NMOS transistors (M3) that together with gain cell transistors M2 implement XNOR functionality. To increase the capacitance at the Q nodes (and consequently improve the retention time), and support the parallel refresh and compare (search), M1 and M2 transistors are implemented as high Vt devices.

The DASH-CAM row, comprising several DASH-CAM cells (32 in our evaluation), ML sense amplifier and precharge circuitry, is shown in Fig. 4(b). The *M*_eval_ transistor shared by all DASH-CAM cells of a DASH-CAM row controls the ML discharge rate by applying different levels of the evaluation voltage *V*_eval_. Since ML discharge rate is used as a measure of the Hamming distance between the query pattern and a dataword stored in a DASH-CAM row, tuning the*V*_eval_ allows user-defined configuration and dynamic adjustment of the Hamming distance threshold.

DASH-CAM has two main operating modes: (1) (approximate) search and (2) refresh. The latter is mandatory to maintain the data stored in dynamic memory cells. To implement these operating modes, and to enable the memory initialization, DASH-CAM supports (1) read, (2) write and (3) compare operations. Read and consecutive write comprise the refresh operation.

DNA basepairs (referred to as *bases* throughout the paper) are encoded using one-hot encoding, for example A is encoded as ‘0001’, G as ‘0010’, C as ‘0100’ and T as ‘1000’. The datawords are stored row by row, as in a typical single-port memory. The write operation is identical to that of the 2T GC-eDRAM. The wordline signal (*W L*) is asserted with V_BOOST_ and the respective dataword is asserted on the bitlines (BL).

The read operation is performed in two phases. In the first phase, the BLs are precharged to half *V*_DD_. In the second phase, the *W L* is asserted, opening M1 transistors of the DASH-CAM cell. As a result, BL voltage level rises slightly above half *V*_DD_ if ‘1’ is stored in the cell, or falls slightly below half *V*_DD_ if ‘0’ is stored in the cell. The column sense amplifiers complete the read operation by comparing the BL voltage with the reference voltage *V*_ref_ = 1/2*V*_DD_. The read operation is destructive and therefore may affect the compare operation when both operations are carried out simultaneously, as discussed further in Section 3.3.

Compare (search) operation is illustrated in Fig. 5. The *inverted* query pattern is asserted on the searchlines (SL), which are separate from the bitlines used for read and write operations, to enable parallel compare and read/write accesses. If the query DNA base in a certain DASH-CAM column matches the stored base (as illustrated in Fig. 5(a)), there is no matchline discharge path through such

**Figure 5:**
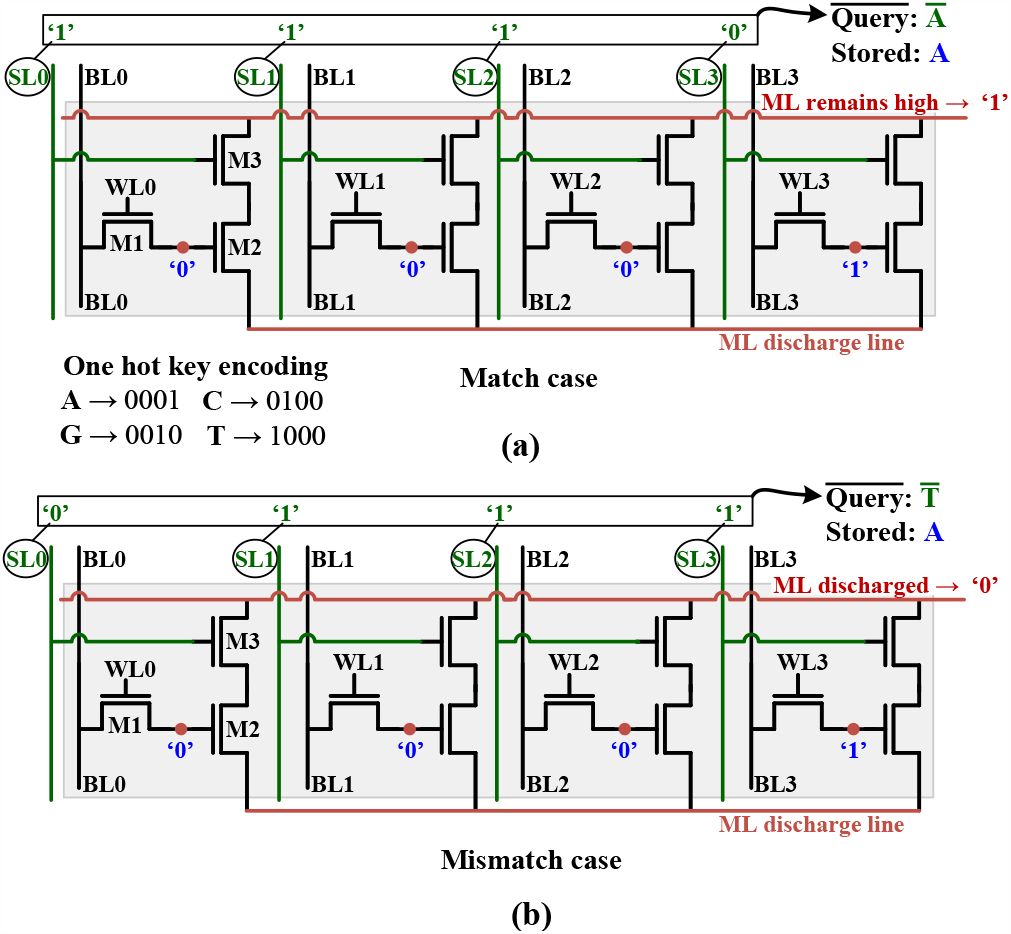
(a) Single-base match in DASH-CAM (b) single-base mismatch in DASH-CAM.

DASH-CAM cell. If however the query base differs from the base stored in the cell (Fig. 5(b)), M2 and M3 transistors in one of the four stacks open, thus enabling a ML discharge path.

One-hot encoding ensures that regardless of what bases are compared (A vs. T, or G vs. C, or T vs. G and so on), the result is always the same: if bases mismatch, then one and only one M2-M3 stack in the cell conducts; when the bases match, all M2-M3 stacks in the cell are shut.

In some cases, individual DNA bases or DNA fragments of either the query pattern or the stored datawords should not affect the result of the compare (i.e., be set as “don’t cares”). To mask off query bases, rendering them “don’t care”, we encode them as ‘0000’. Such combination disables the ML discharge through the cell, which means the comparison results in such cell does not affect the result of such DASH-CAM row. As shown below, this feature is useful vis-à-vis the limited retention time of dynamic storage.

The discharge speed depends on the number of mismatching bases, which define the number of discharging paths. The higher the number of mismatching bases, the higher the ML discharge speed. The Hamming distance tolerance threshold can be dynamically configured by tuning the evaluation voltage *V*_eval_ (Fig. 4(b)), as demonstrated in Section 3.2.

### 3.2 Timing

Fig. 6 shows the timing diagram of DASH-CAM operation comprising two time intervals. In the first interval, a single write command is followed by three consecutive compare (search) commands, each of which takes a single clock cycle divided into two steps: the ML precharge in the first half-cycle and ML evaluation in the second half-cycle. In the precharge step, the SLs are discharged and then the ML is precharged to *V*_DD_ by opening the *M*_PC_ transistor 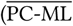 = ‘0’). In the evaluation step, the *M*_PC_ transistor is disabled and the inverted query data is driven onto the SLs. The Hamming distance threshold is set by the evaluation voltage *V*_eval_.

**Figure 6:**
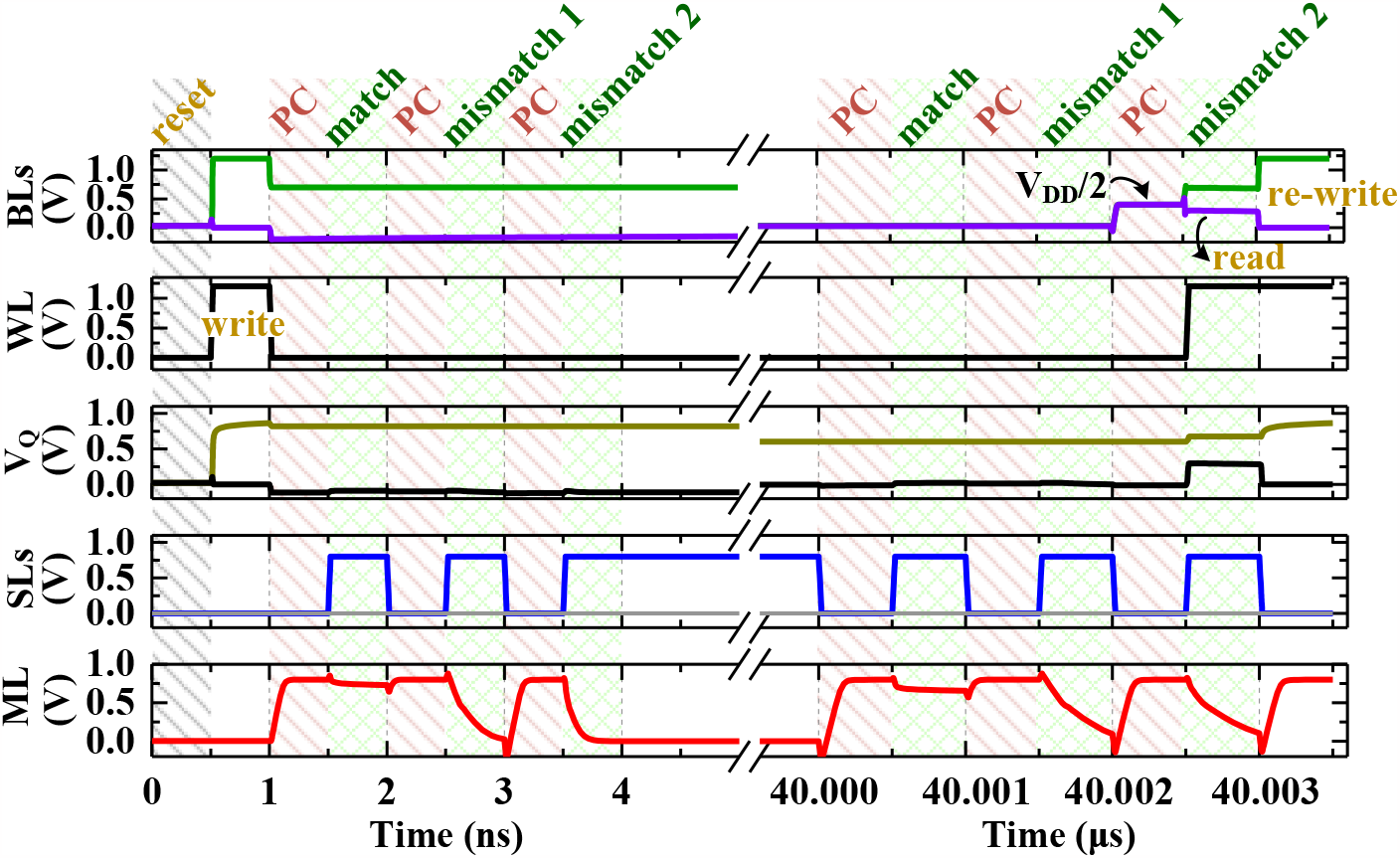
DASH-CAM timing (two intervals).

The ML levels at the end of each cycle signal the compare result: if the ML voltage is above the reference voltage of a sense amplifier (not shown in Fig. 4(b)), the match is signalled by ‘1’ at the output of a sense amplifier; otherwise, a mismatch is signalled by ‘0’ at the output of a sense amplifier.

To enable the exact search operations, *M*_eval_ is fully open (*V*_eval_ = *V*_DD_). The approximate matching operation depends on the conductivity of the *M*_eval_ transistor. During the approximate search, *V*_eval_ is set below the nominal voltage (*V*_eval_ *< V*_DD_), limiting the conductivity of *M*_eval_.

The first compare results in a match while the other two result in mismatches. The Hamming distance leading to the first mismatch is lower than the one in the second, therefore the ML discharges slower during the first mismatch.

In the second interval (which begins at the timestamp of 40*μ*sec), three consecutive compare commands are executed in parallel with a refresh operation shown in the upper part of Fig. 6. It takes one and a half cycles (one cycle for read and half-cycle for write) and is performed in parallel with compare operation. In the first half-cycle, the BLs are precharged to Vdd/2. In the second half-cycle, the WL is asserted. If the cell stores ‘0’, the BL drops slightly below Vdd/2; if the cell stores ‘1’, the BL rises slightly above VDD/2. This difference is detected by column sense amplifiers (not shown). The output of the column sense amplifier is sampled and is asserted on the BL in the third half-cycle, to rewrite the data.

### 3.3 Simultaneous search and refresh

To mitigate the limited retention time of the dynamic storage DASH-CAM is based on, a frequent refresh operation is applied. Since refresh does not change the dataword stored in a DASH-CAM row, it can be done simultaneously with compare (search) operations. Each refresh targets a certain DASH-CAM row and comprises read and write cycles. During read, the dataword is read from a DASH-CAM row. In consecutive write, the same (boosted) dataword is written back. Both read and write operations use wordlines (one at a time) and bitlines. Compare (search) operation uses separate hardware resources (matchlines and searchlines) and therefore is independent of reads and writes, and hence can be performed in parallel.

To analyze the DASH-CAM retention time, we carry out comprehensive Monte Carlo simulations. The resulting retention time distribution is presented in Fig. 7.

**Figure 7:**
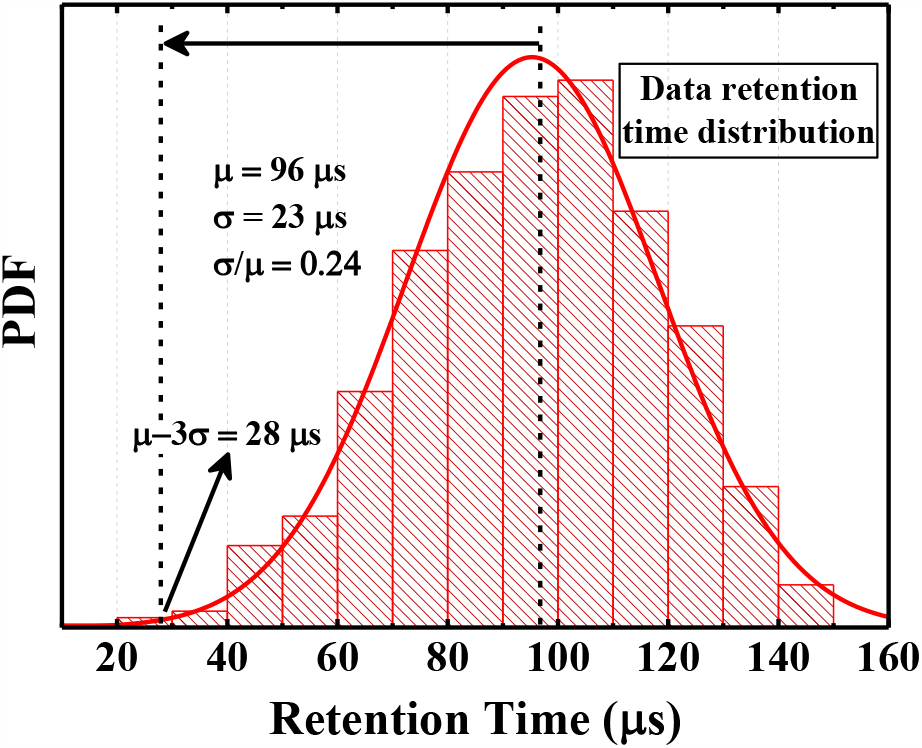
DASH-CAM dynamic storage retention time distribution

Since write only strengthens the charge in dynamic storage, it does not interfere with a simultaneous compare operation (in the same memory row). However, a read from dynamic storage might be destructive and therefore might affect the compare, as follows. Read ‘1’ partially drains the charge from the dynamic storage. In a very unlikely case that the read phase of the refresh occurs in the same DASH-CAM row that is supposed to match during the simultaneous compare, the remaining charge might be falsely identified as ‘0’ (i.e. the parasitic charge at the transistor gate would not be sufficient to open such transistor). Since DASH-CAM cells store one-hot encoded data (i.e. the valid combinations are ‘0001’, ‘0010’, ‘0100’ and ‘1000’), such error will result in a base becoming ‘0000’ (or ‘N’), which is a ‘don’t care’ combination. Hence, such error will not change the true result (a match will not become a mismatch). However, if a very large number of bases are interpreted as ‘N’ (otherwise referred to as *ambiguous’*), it might increase the number of false positives results, where the queries that are supposed to mismatch, match instead because the mismatching bases are masked off by ‘N’. To make sure it does not happen, a compare can be disabled in a refreshed DASH-CAM row. According to our evaluation, disabling a compare in one out of tens of thousands of DASH-CAM rows does not affect its classification accuracy.

During read ‘0’, the bitline charge is shared with a dynamic storage. However, since the (parasitic) capacitance of such dynamic storage is much smaller than that of the entire bitline, the voltage at the gate of the M1 transistor will not reach the bitline voltage of *V*_DD /_2 = 400 mV. Since DASH-CAM cell M1 transistor features the threshold voltage of 420-430mV, it will remain shut, and therefore a false mismatch is quite impossible.

We quantitatively evaluate the limited retention time accuracy impact in Section 4.5.

## 4 APPLICATION & RESULTS

A rapid, accurate, cost-effective and easy to use solution for pathogen detection and identification is critical to enabling the worldwide genomic surveillance and viral pandemic control [6, 42]. In this section, we discuss applying DASH-CAM to the task of viral pathogen classification.

### 4.1 DASH-CAM as pathogen classification

Fig. 8 presents a DASH-CAM-based accelerator for pathogen classification as a part of DNA sequencing and analysis pipeline. The purpose of this design is to classify an unknown sequenced genome into one of *n* predetermined organisms (or creating a notification if such genome does not belong to any of the classes). The predetermined pathogen genomes, each representing a separate class, collectively comprising the reference DNA database, are organized as multiple sets of short DNA fragments, known as *k*-mers, each having a length of *k* basepairs [38]. The reference DNA *k*-mers extraction is illustrated in Fig. 8(b)), where the first *k*-mer is extracted from position 0 to position *k* − 1 of each reference genome, the second *k*-mer is extracted from position 1 to position *k*, and so forth. The *k*-mer extraction stride may vary. The reference DNA database is generated by storing such *k*-mers in the DASH-CAM offline, as shown in Fig. 8(b), where each *k*-mer is stored in a separate DASH-CAM row, and each genome class is stored in an individual DASH-CAM block (a set of DASH-CAM rows, preferably of a size of power of two, to enable an easy identification of each such block by simple address encoding). A typical viral genome comprises several thousands of bases (and consequently *k*-mers). To reduce the memory size, we may select only a fraction of *k*-mers in each reference genome to build the reference database. The impact of such reference genome “decimation” on the classification accuracy is studied in Section 4.4.

**Figure 8:**
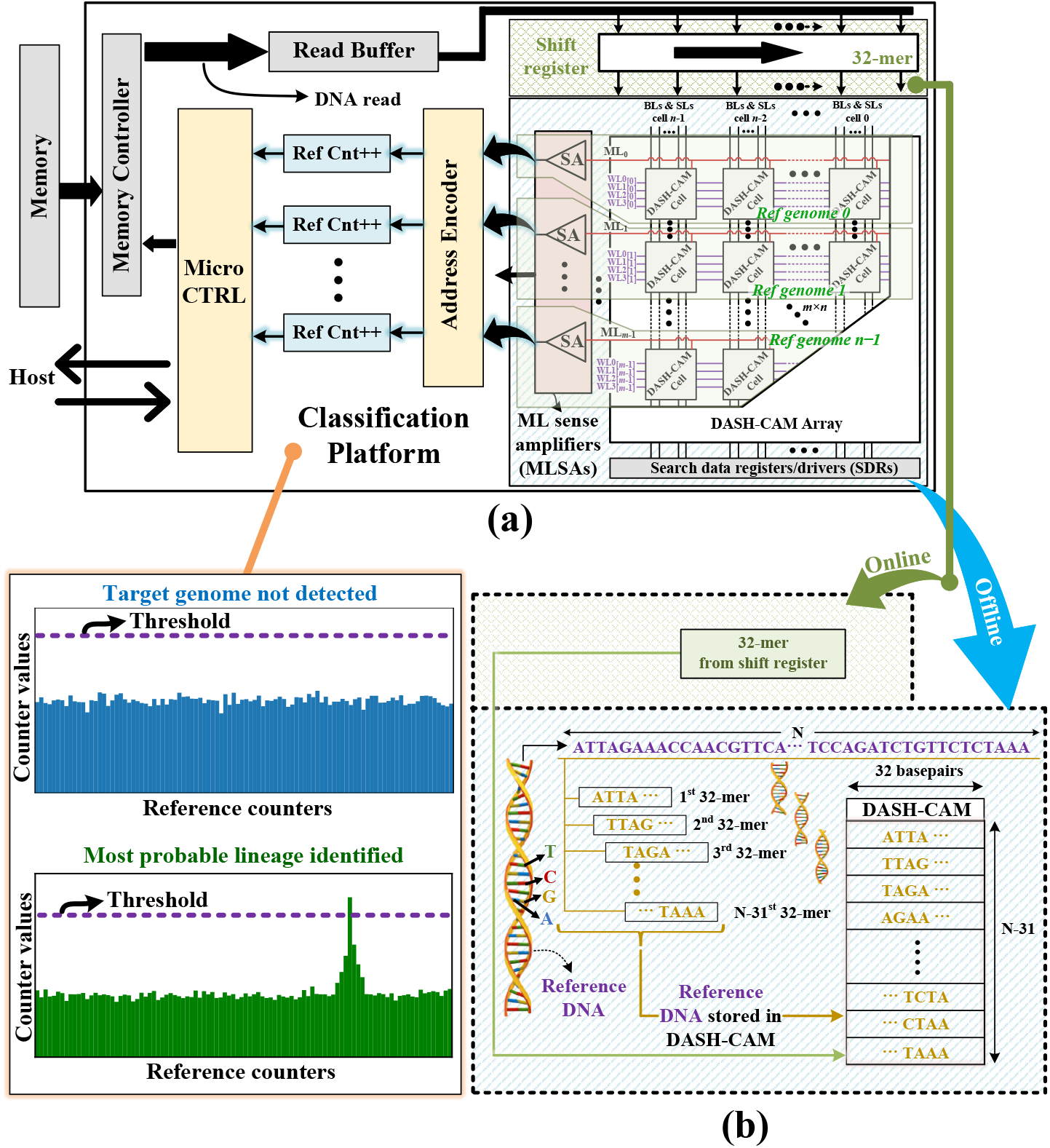
(a) DASH-CAM as a part of a pathogen classification platform: A simplified illustration of DASH-CAM array and the *reference counters* (Ref Cnt); (b) The offline construction of the reference DNA database in DASH-CAM (bottom) and the online classification operation (top). N is the overall length of the reference database. Highlighted in (a): Two possible classification outcomes inferred form the *reference counter* (Ref Cnt) levels’ distribution.

During its operation, DASH-CAM based pathogen classifier (Fig. 8(a)) retrieves the DNA reads from an external memory and transfers them to a read buffer that feeds the shift register. The memory bandwidth required to support the peak DASH-CAM throughput is 16GB /s.

The DASH-CAM array is fed by a 32-base wide segment of the shift register. The DNA read is shifted one base to the right in a sliding window manner in every clock cycle, allowing querying a single 32-mer per cycle. The process is controlled by a microcontroller implemented as a state machine. Its control registers are memory-mapped for accessibility by the host.

In an ideal scenario, if the newly sequenced organism belongs to one of the existing classes, its *k*-mers should match exactly in the DASH-CAM. However, DNA reads are prone to sequencing errors [31] which create variation between the *k*-mers of the newly sequenced genome and the reference DNA. Another source of potential difference are genetic variations, frequent in quickly mutating viral pathogens (such as Severe Acute Respiratory Syndrome coronavirus-2 (SARS-CoV-2)). Such differences result in a nonzero Hamming distance between a *k*-mer and a reference fragment that would otherwise match exactly. The ability of DASH-CAM to tolerate Hamming distance enables accurate classification of DNA reads with multiple sequencing errors or genetic variations. Furthermore, the programmable Hamming distance threshold enabled by DASH-CAM supports a wide variety of sequencing error profiles typical in commercial sequencers [20].

Every time a *k*-mer matches in a certain reference block in DASH-CAM, the *reference counter* associated with such reference block is incremented. Fig. 8(a) illustrates three possible classification outcomes. If by the end of the classification process, no reference counter reaches a certain user-defined configurable threshold, a misclassification notification is generated (signalling that the newly sequenced sample contains no DNA of the target pathogens). If the number of hits exceeds the threshold in one of the counters, the newly sequenced genome is classified into such class.

The DASH-CAM Hamming distance and the configurable classification thresholds can be optimized by training using a validation set, which consists of either simulated DNA reads or DNA reads of known origin (with known classification results). The optimal threshold values that maximize a target criterion, such as *F*_1_ score, can be determined by periodically classifying such validation set and varying *V*_eval_.

### 4.2 Figures of merit for DASH-CAM DNA classification efficiency

Fig. 9 illustrates the mapping of the reference database comprising *n* reference genomes in DASH-CAM, and the examples of true, false and failed-to-place DNA classification results, generated by DASH-CAM. A true positive result (1) occurs when a query *k*-mer of a newly sequenced organism that belongs to one of the classes included in the reference database, matches in a correct position in DASH-CAM. A false negative result (2) is registered when such a *k*-mer fails to match in the right reference class and matches in a wrong class instead. Hence, such false negative result is also considered a false positive for the wrong class such *k*-mer is falsely associated with. False results mainly occur due to sequencing errors and mutations. Another reason has to do with the DASH-CAM design: when too many reference bases become don’t cares (being effectively masked-off) due to dynamic storage loss, more false positive results are possible. A failed-to-place result (3) occurs when such a *k*-mer does not match anywhere in DASH-CAM, which may happen if the reference is not complete (i.e. contains only a part of the genome, in order to save memory space). The impact of such result on the DASH-CAM accuracy is studied in Section 4.4.

**Figure 9:**
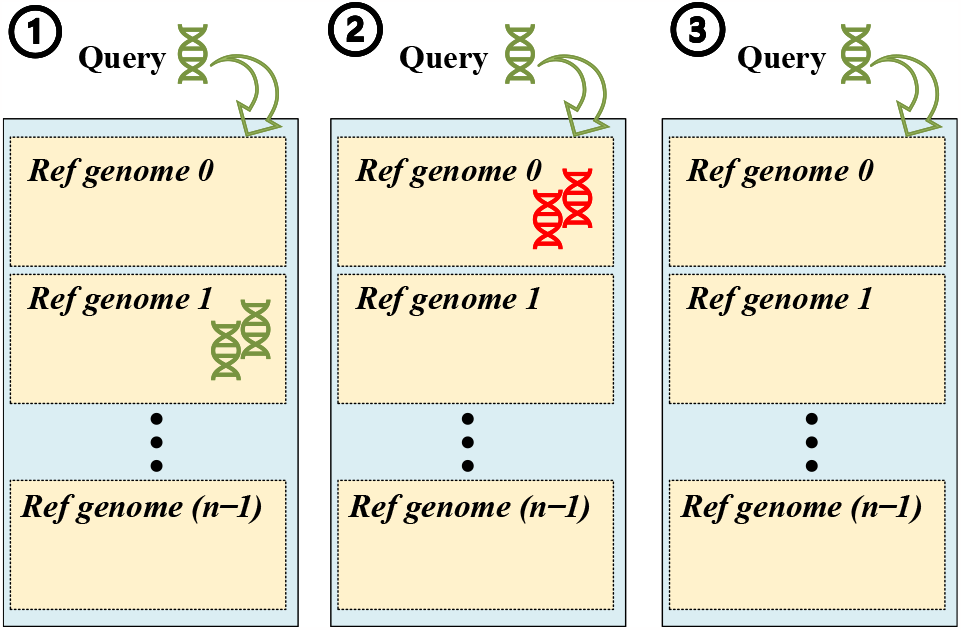
Examples of (1) True, (2) False and (3) Failed-to-place results. In this example, the query *k*-mer be-longs to the organism of reference genome 1.

The figures of merit used in our evaluation are sensitivity and precision, calculated as follows:

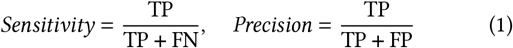

where TP is the number of true positive, FN is the number of false negative and FP is the number of false positive results, respectively.

Additionally, we calculate F_1_ score, which is a harmonic mean of sensitivity and precision. We use state-of-the-art classification tools MetaCahe-GPU [32] and Kraken2 [54] as references for the classification accuracy and performance comparison.

### 4.3 Evaluation methodology and results

DASH-CAM is evaluated using a two-step approach. Its timing, area and power consumption are estimated using extensive Monte-Carlo circuit simulations and full-custom layout, as presented in Section 4.6. DASH-CAM classification efficiency is evaluated using a custom software simulator, as further detailed in this section. The simulated circuit parameters derived at the first step, such as retention time, are used in the classification efficiency evaluation.

All DNA sequences in our evaluation are downloaded from NCBI online data sets [37]. For simplicity, we target relatively short viral genomes, presented in Table 1. We evaluate the DASH-CAM classification accuracy by conducting three separate classification experiments, using virus DNA reads generated by

**Table 1:** Genomes and their sizes used in our study.

| Organism | ~Length (basepairs) |
| --- | --- |
| SARS-CoV-2 | 30,000 |
| Lassa | 10,400 |
| Influenza | 13,500 |
| Rotavirus | 18,550 |
| Measles | 16,000 |
| Candidatus Tremblaya | $\leq 200,000$ |

1. Illumina ART simulator [21],
2. Roche 454 ART simulator [21],
3. PacBioSim simulator [40]; The PacBio reads are generated with 10% error rate.

In all experiments, we attempt to detect and classify the *coronavirus* (SARS-CoV-2), *rotavirus, lassa* virus, *influenza* virus and *measles* virus, as well as *candidatus tremblaya* bacterium (Table 1), in a simulated metagenomic sample, containing DNA reads of the above listed organisms.

To compare DASH-CAM efficiency and performance with those of Kraken2, a Kraken2 database containing the DNA of organisms of Table 1, was created. Similarly, MetaCache-GPU minhashing sketch representation was created. Both tools were applied to our simulated metagenomic dataset, with the *k*-mer size of 32.

The sensitivity, precision and F_1_ score (as the functions of the Hamming distance threshold) for all experiments are presented in Fig. 10(a)-(i). The classification efficiency figures of Kraken2 and MetaCache-GPU do not depend on DASH-CAM Hamming distance threshold; hence, their sensitivity, precision and F_1_ score are presented by horizontal lines. We observe that DASH-CAM sensitivity grows (due to the decreasing number of false negative results) with the increasing Hamming distance threshold for the Roche 454 and PacBio reads (DASH-CAM sensitivity when classifying Illumina reads is 100% due to the high accuracy of such reads). At the same time, DASH-CAM precision diminishes (due to the growing number of false positive matches) in all runs. The precision never reaches zero because it is bounded by the ratio of the number of query *k*-mers of the target species to the number of query *k*-mers of the rest of the species in the metagenomic sample.

**Figure 10:**
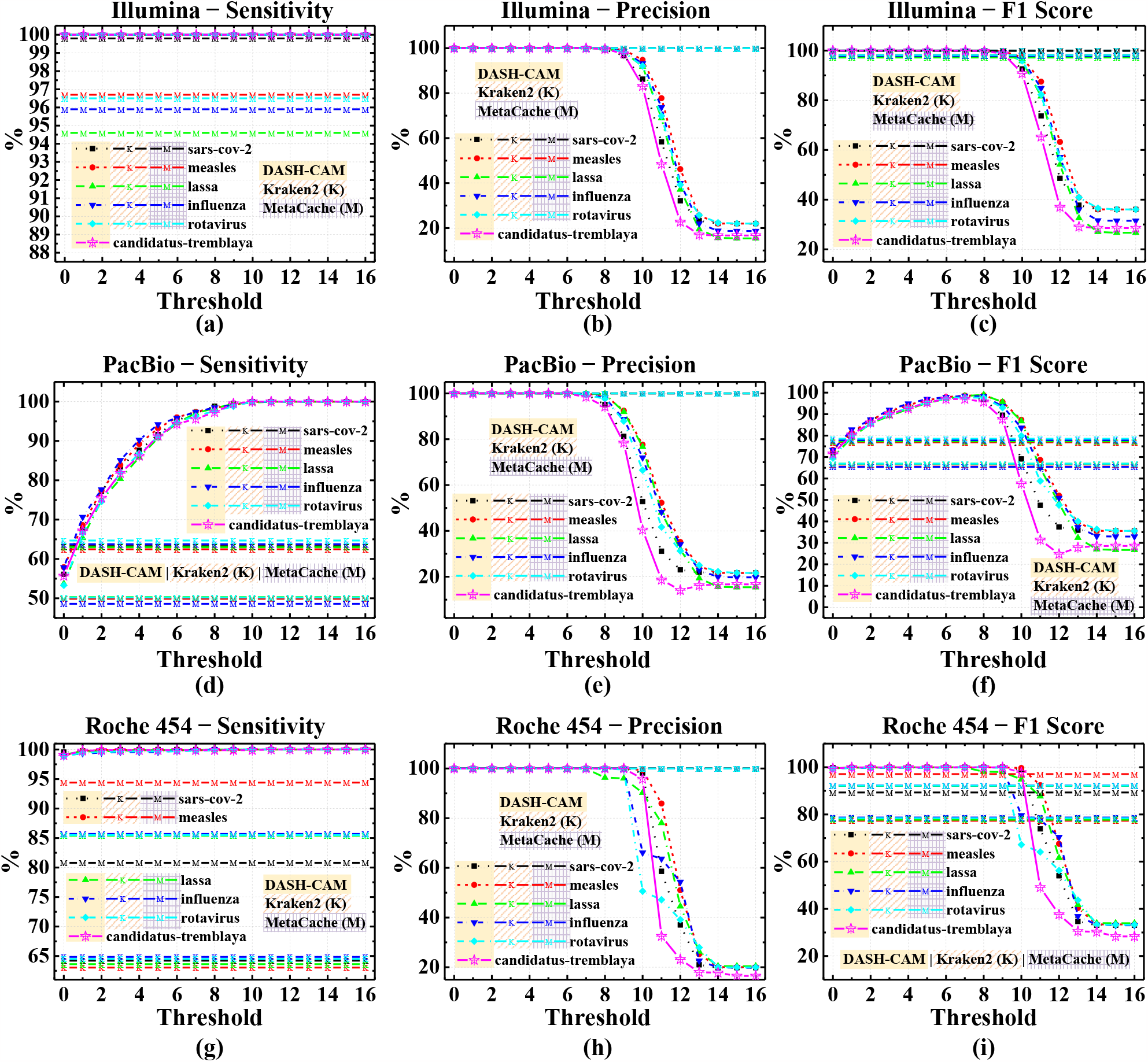
DASH-CAM vs. Kraken2 and MetaCache-GPU: Sensitivity, Precision and *F*_1_ score for (a–c) Illumina reads, (d–f) PacBio reads with 10% error rate (g–i) Roche 454 reads

For high precision Illumina reads, the best F_1_ score is achieved at Hamming distance threshold of 0 (exact match). However, for Roche 454 as well as the higher error rate PacBio reads, the behavior is different. In both these cases, there is a Hamming distance threshold at which the drop in precision begins to outpace the growth in sensitivity. This marks the optimal F_1_ score which occurs in the Hamming distance threshold region of 1-5 (depending on the organism) for Roche 454 reads and 8-9 for PacBio 10% error rate reads respectively.

Based on the results of these experiments, we conclude as follows:

1. F_1_ score has an optimum region. For the low Hamming distance threshold values, the growing DASH-CAM sensitivity outpaces the decreasing precision, leading to the F_1_ score growth. However, as the Hamming distance threshold increases, the drop in precision surpasses the rise in sensitivity, causing the F_1_ score to diminish.
2. The Hamming distance threshold for the optimum F_1_ score varies for different sequencers as a function of DNA reads’ precision. The lower the sequencing error rate, the lower the optimal Hamming distance threshold.
3. DASH-CAM outperforms both Kraken2 and MetaCache-GPU when classifying erroneous DNA reads.

### 4.4 The impact of the reference size on DASH-CAM accuracy

Saving complete reference genomes in DASH-CAM leads to a large memory footprint, which in turn adversely affects DASH-CAM power consumption and silicon cost. Additionally, storing the entire reference genome for each class would require supporting reference blocks of different sizes (as presented in Table 1). Limiting each reference genome to a small reference block of a fixed size mitigates these problems but, in turn, results in occasional failures-to-place as presented in Section 4.2.

In this section, we discuss the impact of the reference block size on DASH-CAM classification accuracy. The reference dataset is created by randomly extracting several thousand *k*-mers (k=32) from each reference genome class. The query set is composed of the same DNA reads, including also the *k*-mers that are not present in the reduced reference dataset.

Fig. 11 shows the resulting *F*_1_ score as a function of the reference block size for Hamming distance thresholds of 0, 4 and 8. For high quality Illumina reads, *F*_1_ score is slightly lower for a small reference block size of 1,000 *k*-mers (for example, for SARS-CoV-2, the *F*_1_ score is 92%, whereas 1,000 *k*-mers holds only 3% of the full reference). However, as the reference size grows (to 6,000 *k*-mers, which is approximately 20% of the SARS-CoV-2 reference size), *F*_1_ score reaches 100%, as with the complete reference.

**Figure 11:**
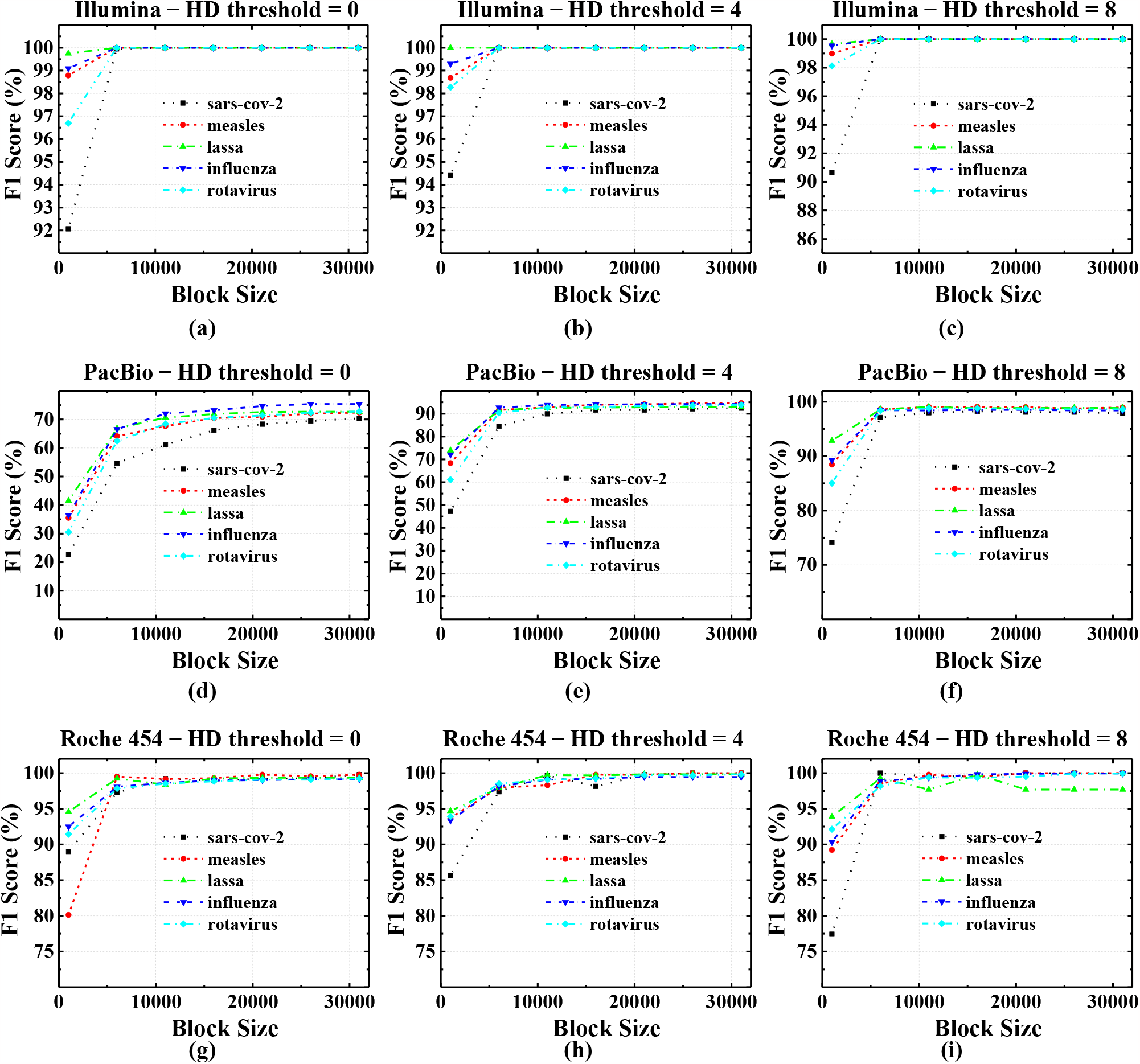
The effect of the reference size on *F*_1_ score for Hamming distance (HD) thresholds of 0, 4 and 8, for (a–c) Illumina reads, (d–f) PacBio reads with 10% error rate (g–i) Roche 454 reads.

The erroneous PacBio reads exhibit a similar behavior, with *F*_1_ growing quickly with the reference size. However, *F*_1_ for PacBio reads is strongly affected by the Hamming distance threshold: for example, for the reference block size of 1,000 *k*-mers, *F*_1_ is 23% for SARS-CoV-2 for the threshold of 0, growing to 74% for the threshold of 8.

To summarize: while DASH-CAM classification accuracy is adversely affected if the reference size is very small (3% for SARS-CoV-2), it quickly reaches its maximum level when the reference block size stands at 20%-40% of the complete reference (20% for SARS-CoV-2).

### 4.5 The impact of data retention time variation on DASH-CAM accuracy

Dynamic storage is prone to retention time variation. Specifically, some of the DASH-CAM cells may lose their charge (due to leakage) sooner than others. In worst cases, such charge loss may occur more frequently than the refresh period. The purpose of this study is to evaluate the impact of retention time variation on DASH-CAM accuracy.

DASH-CAM stores DNA bases in one-hot code (A is encoded as ‘0001’, G as ‘0010’, C as ‘0100’ and T as ‘1000’). A charge loss manifests in ‘1’ becoming ‘0’, i.e. a one-hot codeword turns into ‘0000’. Such code effectively masks off the base (in other words, ‘0000’ is a don’t care, which does not affect the results of the compare operation, because four zeros cut the matchline discharge path through that cell regardless of the query base value).

To evaluate the retention time variation, we model the charge in a DASH-CAM cell as an exponentially decaying function *e* ^−*t*/τ^ where τ is a random variable distributed close to normally as shown in Fig. 7.

Fig. 12 shows the sensitivity and precision as functions of time, for all organisms presented in Table 1, for low-quality PacBio reads with Hamming distance threshold of 0. While for approximately 95*μ*s, the precision is at 100%, the sensitivity grows due to the decreasing number of false negative results (following the growing fraction of bases being masked off). Between 95*μ*s and 102*μ*s, the precision drops to its lower bound (because the false positive results quickly reach maximum) while the sensitivity hits 100%.

**Figure 12:**
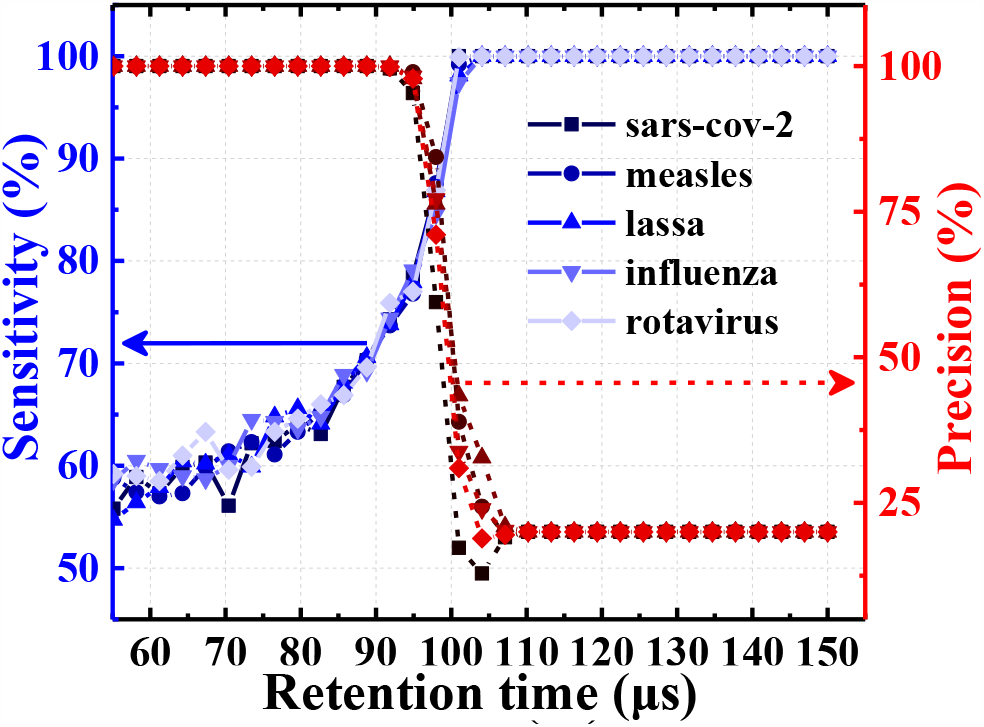
DASH-CAM sensitivity and precision as function of time for PacBio reads with 10% error rate.

Following this study, we set the refresh period at 50*μ*sec, which allows refreshing the entire reference (assuming that all reference blocks are refreshed separately and in parallel), while being sufficient to keep the probability of retention time-related classification accuracy loss close to zero.

### 4.6 Power consumption, silicon area and speedup

We designed DASH-CAM using a commercial 16nm FinFET process and used post-layout results to evaluate its timing, area and power consumption. The DASH-CAM 12T cell area shown in Fig. 13 is 0.68 μm^2^. DASH-CAM operates at 700 mV and consumes an average of 13.5fJ per 32-cell row.

**Figure 13:**
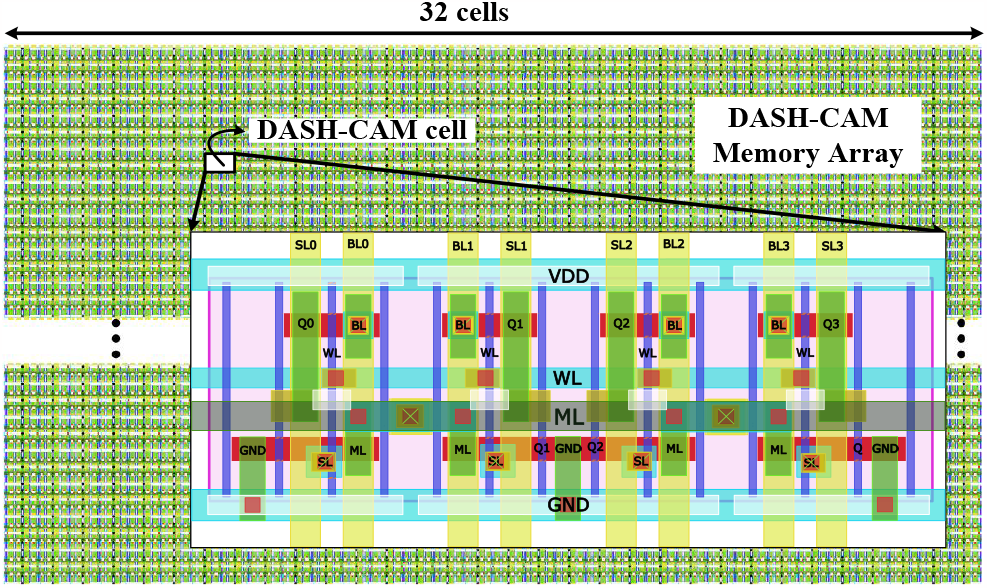
DASH-CAM array layout. Highlighted: DASH-CAM 12T cell

These results are summarized in Table 2, and compared with prior art designs for *k*-mer or pattern matching: HD-CAM [15], EDAM [20] and 1R3T resistive TCAM [10]. The main advantage of DASH-CAM over HD-CAM and EDAM is its density, which enables efficient classification of larger genomes, such as bacterial pathogens. The advantages compared to 1R3T resistive TCAM are the unlimited write endurance, as well as the ability to perform approximate search and tolerate large Hamming distances.

**Table 2:** DASH-CAM area, power, and latency, compared with HD-CAM [15],EDAM [20] and 1R3T resistive TCAM [10].

| Design | DASH-CAM | HD-CAM | EDAM | 1R3T TCAM |
| --- | --- | --- | --- | --- |
| Technology | 16 nm | 65 nm | 65 nm | 90 nm |
| Cell type | 12T | 30T | 42T | 1R3T |
| Cell area | $0.68\mu m^2$ | $5.45\mu m^2$ | $33\mu m^2$ | $4 \times 1.57\mu m^2$ |
| Search time | 1 ns | 1 ns | 0.75 ns | 0.96 ns |
| Search energy/cell | 0.3 fJ | 0.54 fJ | 1.4 fJ | $4 \times 0.51$ fJ |

Assuming the reference block size of 10,000 *k*-mers, the DASH-CAM that can classify viral genomes into 10 classes of concern, has the area of 2.4 sq mm, and consumes 1.35W.

Based on timing results of extensive Monte-Carlo simulations, DASH-CAM can be operated at 1GHz. Since refresh is performed in parallel with search operation, it does not affect the overall DASH-CAM performance.

We compare DASH-CAM with Kraken2 and MetaCache-GPU. We use Intel XEON server with 48 cores at 2.2GHz and 380 GB DDR4, and NVIDIA RTX A5000 GPU (8192 CUDA cores, 256 Tensor cores, 64 RT cores), with 24 GB GDDR6 Memory and up to 768 GB/s DRAM bandwidth. The average MetaCache-GPU and Kraken2 classification throughput figures are 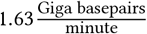 (Gbpm) and 1.84Gbpm, respectively.

DASH-CAM processes one *k*-mer per cycle, hence its classification throughput is *f*_*op*_ × *k*, where *f*_*op*_ is the DASH-CAM operating frequency and *k* is the *k*-mer size. For *f*_*op*_ = 1 GHz and *k* = 32, the classification throughput is 1, 920 Gbpm, achieving an average speedup of 1, 040× and 1, 178× over Kraken2 and MetaCache-GPU respectively.

## 5 CONCLUSION

In this paper, we present DASH-CAM, a novel dynamic storage based approximate search-capable content addressable memory, designed for computational genomics applications. DASH-CAM enables parallel search with a user configurable Hamming distance tolerance. DASH-CAM cell features 12 transistors, reaching 5.5× higher memory density compared with state of the art static solutions. The Hamming distance tolerance is defined by the speed of matchline discharge during the approximate search. A special perrow transistor is used to set a user-configurable Hamming distance threshold. We apply DASH-CAM to the task of rapid and accurate detection and classification of viruses of epidemic significance, thus enabling an efficient pathogen transmission tracking and genomic surveillance. We show that depending on the sequencing error rate, DASH-CAM provides up to 30% and 20% higher *F*_1_ score compared to state-of-the-art DNA classifiers MetaCache-GPU and Kraken2, while outperforming them by 1,178× and 1,040×, respectively.

## ACKNOWLEDGMENTS

This work was supported by the European Union’s Horizon Europe programme for research and innovation under grant agreement No. 101047160. The work of Esteban Garzón was supported by the Italian MUR under the call “Horizon Europe 2021-2027 programme – H25F21001420001”

## Notes

### Competing Interest Statement

The authors have declared no competing interest.

